# Metabolic PCTA-Based Shift Reagents for the Selective Detection of Extracellular Lactate Using CEST MRI

**DOI:** 10.1101/2024.07.09.602789

**Authors:** Remy Chiaffarelli, Pedro F. Cruz, Max Zimmermann, Carlos F. G. C. Geraldes, Paul Jurek, André F. Martins

## Abstract

Lactate is a key metabolic driver in oncology and immunology. Even in the presence of physiological oxygen levels, most cancer cells upregulate anaerobic glycolysis, resulting in abnormal lactate production and accumulation in the tumor microenvironment. The development of more effective, sensitive, and safe probes for detecting extracellular lactate holds the potential to impact cancer metabolic profiling and staging significantly. Macrocyclic-based PARACEST agents have been reported to act as shift reagents (SRs) and detect extracellular lactate via CEST MRI. Here, we introduce a new family of SRs based on the PCTA ligand, an inherently stable and kinetically inert group of molecules with the potential for (pre)clinical translation. We observed that Yb-PCTA and Eu-PCTA can significantly shift lactate -OH signals in CEST spectra. *In vitro*, CEST MRI experiments proved that imaging extracellular lactate with these complexes is feasible and maintains high specificity even in the presence of competing small metabolites in blood and the tumor microenvironment. *In vivo*, preclinical imaging demonstrated that Yb-PCTA can be safely administered intravenously in mice to detect extracellular lactate non-invasively. This work represents a significant step towards precision and molecular imaging, demonstrating that the PCTA-ligand is a promising scaffold for developing molecular imaging sensors. These chemical sensors have broad medical applications, particularly in oncology and physiology.

---

Metabolic imaging, the non-invasive visualization of metabolite distribution within living organisms, is essential for understanding disease processes and assessing treatment responses. It enables the early detection, characterization, and monitoring of relevant diseases, e.g., in oncology and neurology, and therefore holds paramount significance in medicine and biomedical research. A key aspect of metabolic imaging is that it allows monitoring of the dynamic behavior of cells and tissues, offering the potential to identify abnormal metabolic patterns associated with disease states. For example, cancer cells overproduce lactate even in the presence of high oxygen and glucose levels, a phenomenon called the Warburg effect. Most of the lactate produced by cancer cells builds up in the extracellular environment, affecting the tumor and the homeostasis of the surrounding tissues.^1^ Detecting the lactate build-up in tissues is thus critical as it translates to a precise measure of cancer metabolic activity and may provide a direct readout of cancer staging. Modern imaging technologies offer substantial promise for assessing cancer metabolic activity and tumor staging. Magnetic resonance spectroscopy, aided by hyperpolarized ^13^Clabelled substrates like pyruvate^2^ and fumarate^3^ or deuterated metabolites like glucose and acetate,^4^ provides invaluable information regarding enzymatic activity in tumors related to staging and therapy response. These methods, however, lack spatial resolution and cannot distinguish between intra- and extracellular metabolic processes, preventing accurate quantification of extracellular lactate produced by the tissues. The chemical shift between lactate hydroxyl and water protons allows the detection of lactate with Chemical Exchange Saturation Transfer (CEST) techniques;^5,6^ however, it does not aid in the discrimination between extra- and intracellular lactate. The water signal obscures the slight chemical shift of free lactate (<1 ppm), hampering the quantification of small lactate concentrations. Hence, a method for non-invasively imaging extracellular lactate produced by the cancer cells is of utmost importance to better understand metabolic dynamics and compartmentalization.

Recently, a family of shift reagents (SRs) has demonstrated remarkable capabilities in selectively detecting extracellular lactate using CEST. These CEST SRs consist of stable inorganic complexes formed by the complexation of lanthanide ions with DOTA-type macrocycles. A Yb^3+^ complex based on a derivative of DO3A, Yb-MBDO3AM, was reported to bind to lactate at different concentrations.^7^ A simplified version of these SRs utilized the Eu-DO3A complex, which coordinates with lactate by displacing the two water molecules directly bound to the inner sphere of the lanthanide.^8^ This specific interaction creates a ternary complex with lactate, allowing for the precise detection of the lactate -OH group, effectively shifting it away from the bulk water signal. When co-administered in the ternary complex with lactate, Eu-DO3A could readily detect extracellular lactate excreted in the bladder of a healthy mouse. Despite detecting the intact Eu-DO3A form in the urine, lanthanide complexes formed with DO3A show a lower kinetic inertness than most octacoordinated Ln-DOTA complexes. Kinetic inertness is a crucial parameter when designing medical imaging inorganic contrast agents directed to clinical translational applications, as the release of lanthanide ions (e.g., Gd^3+^) may lead to severe physiological complications associated with nephrogenic systemic fibrosis (NSF) in patients with renal impairment.^9^ In this pursuit, the pyclen-based PCTA ligand proved to be an important alternative to cyclenbased DOTA-type macrocycles due to its high thermodynamic stability (log K_ML_ = 18.15-20.63 for Ce-Yb) and kinetic inertness (T_1/2_ in the range of hours) similar to Ln-DOTA complexes.^10^ Like Ln-DO3A complexes, Ln-PCTA complexes generally show higher hydration (q =2) and faster water exchange rates (*k*_*ex*_) than Ln-DOTA complexes due to their increased rigidity akin to Ln-DO3A.^11–15^ The heptadentate PCTA ligand forms Ln^3+^ complexes highly suitable for safe translational applications. Despite the increased hydration of Ln-PCTA complexes, which allows other biogenic molecules to replace the water molecules in the coordination sphere. In 2022, the FDA approved Gd-PCTA (gadopiclenol) as a new MRI contrast agent. It is considered more effective than traditional Gd-based MRI contrast agents with octadentate ligands.^16^

Here, we describe the development of a new generation of SRs based on Ln-PCTA for selective detection of extracellular lactate by CEST MRI. For this purpose, we selected Ln^3+^ ions that have a minor impact on the *T*_1_ relaxation of water protons in the concentration ranges relevant for CEST MRI: Eu^3+^, Yb^3+^, and Pr^3+^.^17^ The interaction between Ln^3+^-based PCTA complexes (Yb-PCTA, Eu-PCTA, and Pr-PCTA) and lactate was characterized by using high-resolution Nuclear Magnetic Resonance (NMR), revealing the formation of lactate-Ln-PCTA ternary complexes. The CEST effect of these complexes was quantified *in vitro* at 7T, demonstrating a specific signal for lactate. Additionally, binding affinity values indicated a weak interaction, suggesting that lactate was not significantly depleted from the bloodstream when these SRs were used for *in vivo* imaging. *In vitro* and cytotoxicity assessments validated the safety and feasibility of SRs for *in vivo* imaging. A preclinical animal study using Yb-PCTA showed safe detection of the lactate-Yb-PCTA ternary complex in the bladder, emphasizing the translational potential of this approach. This work highlights the capability of lanthanide-based PCTA complexes and CEST MRI for metabolic imaging applications, offering a new tool for profiling various cancer cell lines and the associated tumor microenvironment.

## Results and Discussion

In this study, we investigated whether a new generation of SRs based on Ln-PCTA was able to detect extracellular lactate and provide selective detection of extracellular lactate by CEST MRI *in vitro* and preclinically. We prepared several solutions containing Lncomplexes (Ln: Eu^3+^, Yb^3+^, and Pr^3+^) to explore the potential formation of ternary complexes between Ln-PCTA and lactate. Phantom imaging experiments were performed in serum to mimic physiological conditions and at pH 6 and 7 to more closely resemble acidic conditions often found in the tumor microenvironment, where lactate accumulates.^18^ Eu^3+^, Yb^3+^, and Pr^3+^ were selected based on the expected induced chemical shifts resulting from their large magnetic moments,^19^ and their limited impact on the longitudinal and transversal relaxation of water protons (*r*_1_, *r*_2_) compared to other Ln^3+^ such as Gd^3+^. The PCTA complexes employed in the study were synthesized as described in the supplementary information (see S.I.). As depicted in Figure 1, a ternary complex can be formed between the Ln-PCTA complexes and lactate by replacing at least one of its two inner-sphere water molecules, as observed before between Ln-DO3A-type complexes with lactate and other α-hydroxy-carboxylates. ^8,20^ In an aqueous solution, DOTA-type complexes show the presence of twisted square antiprismatic (TSAP) and square antiprismatic (SAP) diastereoisomeric pairs of enantiomers, which result from the combination of the two square [3,3,3,3] conformations of the cyclen macrocyclic ring, in which the four ethylenediamine N-C-C-N bridges adopt the identical gauche conformations of opposite helicities, (δδδδ) or (λλλλ), with the two possible orientations of the acetate arms attached to the macrocyclic nitrogen atoms, Λ (clockwise) and Δ (counterclockwise).^21^ Eu- and Yb-DO3A were reported to show a preferred SAP structure, with the metal-coordinated lactate having the OH group in an apical position and showing highly shifted CEST peaks of approx. 45 and 150 ppm, respectively.^8,20^ A less favored TSAP structure produced a smaller shift (16 ppm) for the CEST peak of Eu-DO3A-lactate with the OH group in a more equatorial position.^8^ However, in the case of Ln-PCTA, the rigidity of the pyridine group in the pyclen ring makes it impossible to adopt the [3,3,3,3] conformation. Instead, it adopts a [4,2,4,2] conformation bisected by a mirror plane, with the nitrogen atoms located in the center of each side of the ring and the three ethylenediamine N-C-C-N bridges having (δλδλ) conformations. The Ln-PCTA complexes feature three acetate arms and can adopt twisted snub disphenoid (TSD) coordination geometries. A 3D structure in Figure 2B shows the proposed interaction between Ln-PCTAs and lactate. These geometries exhibit both (δλδλ)Λ and (δλδλ)Δ orientations, and due to the rigidity of the pyridine unit, the four nitrogen atoms are not coplanar. Consequently, the TSAP and SAP conformations typical of DOTA-like complexes are unattainable.^22,23^ Taking this information into account, we determined the diastereoisomers of Ln-PCTA-lactate in solution using high-resolution ^1^H NMR spectra of Yb-, Eu- and Pr-PCTA in the absence/presence of two equivalents of lactate, at various temperatures and pH values. The Ln-PCTA complexes are highly fluxional at high temperatures, with a fast exchange in the NMR time scale at 90ºC, between the dominant TSD (δλδλ)Λ isomer (10 resonances, as expected from the structure) with the very low populated (< 10%) TSD (δλδλ)Δ isomer, leading to extensive broadening of most resonances at 25ºC.^10^ In the presence of lactate, the ^1^H NMR spectra at 90ºC showed two extra signals (2.01 and 5.42 ppm for Yb-PCTA-lactate) corresponding to the CH^3^ and CH groups of coordinated lactate in the Ln-PCTA-lactate ternary complexes in solution, which displayed only the TSD (δλδλ)Λ isomer. Again, this isomer is in fast exchange with the very minor (δλδλ)Δ isomer, as reflected in an extensive broadening of most resonances at lower temperatures (Fig. 2A, S4-6) and the presence of exchange cross-peaks in the EXSY spectrum of Eu-PCTA different from those of the COSY spectrum (Figure S7).^20^ The resonances of bound lactate exhibit some broadening at 90ºC, compared to the free lactate resonances. However, as temperatures decrease, they gradually broaden until disappearing entirely at ∼0ºC, suggesting a fast-to-intermediate exchange regime in their binding to Ln-PCTA (Fig. S5-6). This suggests a relatively weak interaction between lactate and the Ln-PCTA complexes, a finding supported by CEST titrations (Table 1, Fig. S2). This weak interaction may be attributed to the TSD conformation of the complex, where the four nitrogen atoms of the pyclen ring do not align in the same plane—two are positioned above and two below the median plane. Consequently, the three Ln-bound carboxylate oxygens are located slightly off the single plane. This distortion of the two available lactate-binding positions within Ln-PCTA could lead to steric hindrance in lactate binding. Additionally, a recent report shows how a 100-fold excess of lactate induces a 25% decrease of the *r*_1_ of Gd-PCTA, confirming the meager binding.^24^ While the axial CH_2_ protons from the macrocyclic ring in PCTA complexes showed less shifted resonances compared to the corresponding DO3A complexes -typical of TSAP structures, we observed shifts for all the proton resonances and broadening of some of them when lactate was bound. Hence, we anticipated a reasonable -OH CEST shift in the CEST spectra for the lactate -OH group as it exchanges with the surrounding bulk water. However, for efficient CEST imaging, the *T*_1_ relaxation times of the exchanging system must be sufficiently long to ensure that the CEST signal does not decrease rapidly during the acquisition.^25^ We have determined *T*_1_ values and the corresponding *r*_1_ relaxivity, a key governing parameter for CEST detection, in blood serum at 37°C. We measured *r*_1_ values of 0.009 mM^-1^·s^-1^ for Yb-PCTA, 0.003 mM^-1^·s^-1^ for Eu-PCTA, and 0.007 for mM^-1^·s^-1^ Pr-PCTA. Therefore, the impact of longitudinal relaxation on CEST detection can be considered negligible.

**Table 1.** Binding and exchange rates of lactate·SRs.

| | pH | $K_A$ ($M^{-1}$) | $k_{ex}$ ($s^{-1}$) |
| --- | --- | --- | --- |
| Yb-PCTA | 6 | 20 | 1270 |
|  | 7 | 11 | 1770 |
| Eu-PCTA | 6 | 100 | 1810 |
|  | 7 | 25 | 2820 |
| Yb-DO3A | 6.5 | N/A | 1900 [a] |
| Eu-DO3A | 6 | 45 [c] | 2470 [b] |
|  | 7 | 42 [c] | 2180 [b] |

Binding affinities ( $K_A$ ) and exchange rates ( $k_{ex}$ ) for Yb- and Eu-PCTA were determined as described in S.I. [a] Determined at 298 K at 9.4 T by titrating $B_1$ from 2.34 to 23.4 μT by Zhang et al. 2017. [b] Determined at 298 K at 9.4 T by titrating $B_1$ from 2.35 to 23.5 μT by Zhang et al. 2017. [c] Determined by fitting CEST titrations collected using a $B_1$ of 23.5 μT by Zhang et al. 2017.

**Figure 1.**
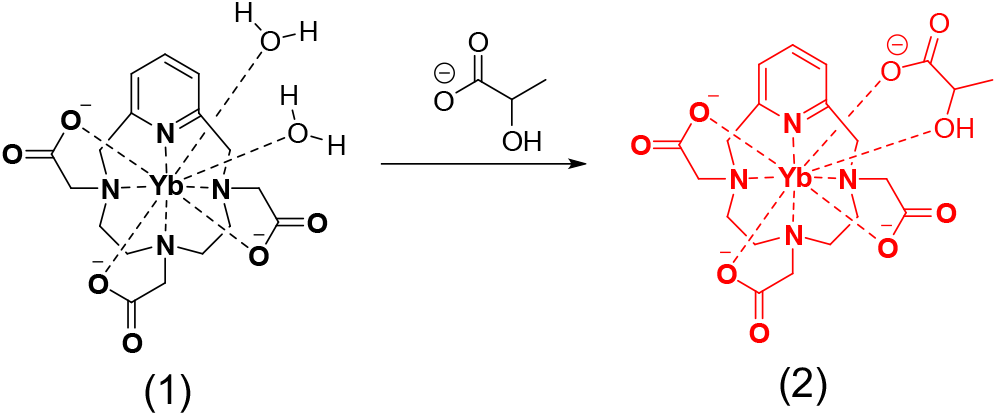
Illustrative schematic of the formation of a ternary complex lactate-Yb-PCTA.

**Figure 2.**
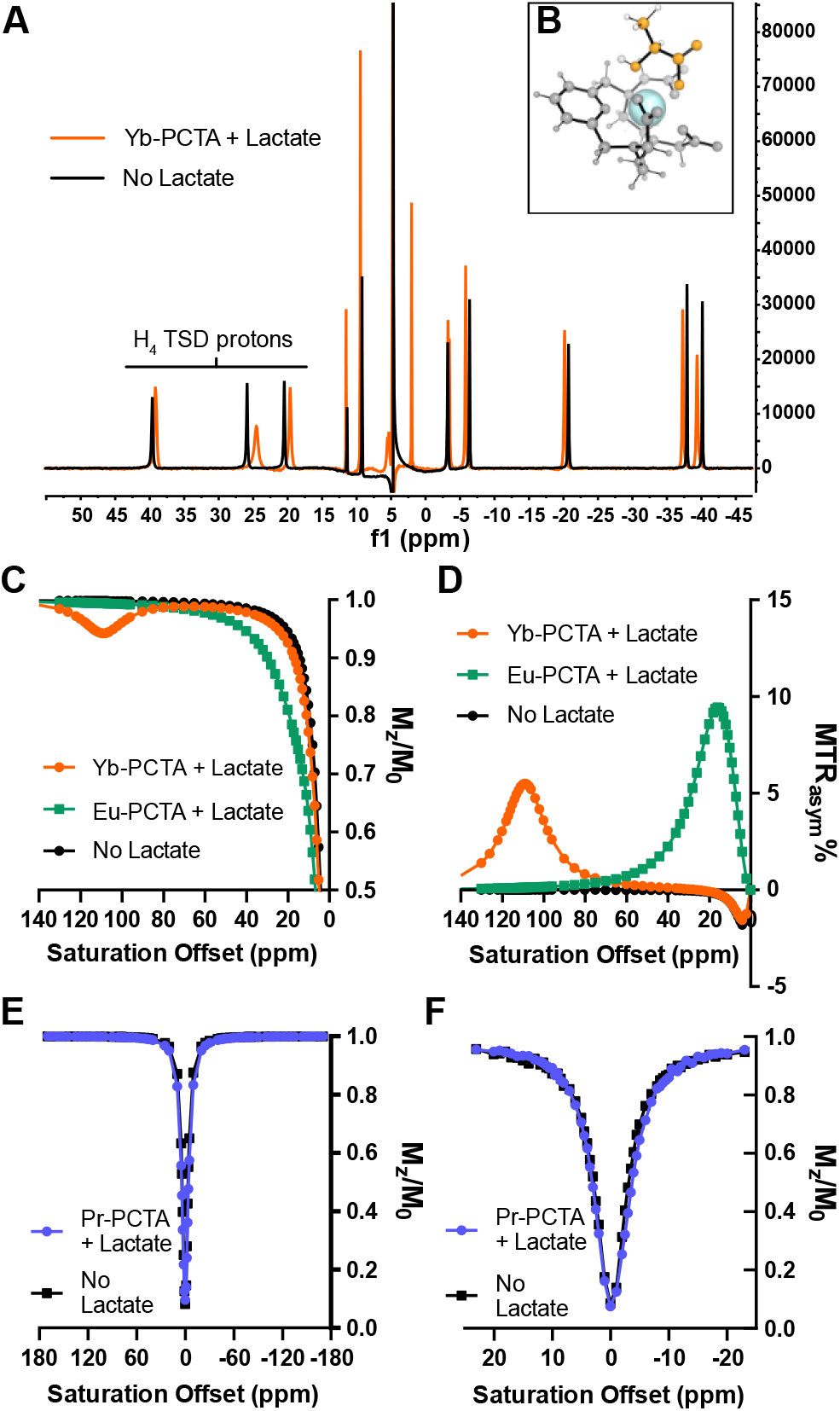
**A**. ^1^H NMR spectra of a solution containing 3 mM Yb-PCTA (black line) and 2:1 Lactate-Yb-PCTA mixture (orange line), acquired at 400 MHz, 363 K, pH 7.1. **B**. 3D schematic of the interaction lactate-Ln-PCTA. **C, D**. z-spectra (C) and MTRa-sym% (D) plot of 50 mM Yb- and Eu-PCTA in HEPES buffer with 50 mM lactate, acquired at 7T MR after pre-saturation pulses of 16 µT and 5 s, at 298 K, pH 6.7. “no lactate”: spectrum of Eu-PCTA without lactate. CEST spectra were fitted to Lorentzian-line shapes using a two-pool model (lactate-Ln-PCTA and water). **E**. z-spectra of 1:1 20 mM lactate-Pr-PCTA or 20 mM lactate, acquired with 5 s 16 µT saturation pulses, pH 6. **F**. z-spectra of 1:1 40 mM lactate-Pr-PCTA or 40 mM lactate, acquired in a narrower spectral bandwidth with 5 s 20 µT saturation pulses, pH 6.

We performed CEST MRI experiments in phantoms containing one of the three Ln-PCTA complexes in the absence and presence of lactate at different concentrations. CEST signals originating from lactate were readily detected using a 7T MR scanner for Yb and Eu complexes (Fig. 2C-D). However, Pr-PCTA (20 and 40 mM) did not show a significant CEST effect in the presence of lactate (Fig. 2E-F). In the presence of one lactate equivalent, Yb-PCTA produces a prominent and distinct CEST peak at around 109 ppm, compared to the typical 0.6-0.8 ppm for non-coordinated lactate. Meanwhile, the shift in the CEST signal induced by Eu-PCTA was slightly smaller than that of the previously reported with Eu-DO3A (approximately 14 ppm). The effect is even more evident when a lower saturation power (8 µT) is used thanks to a sharper definition of the water peak and reduced magnetization transfer (MT) effects (Fig. S1). The quantification of the CEST effect, represented by the Magnetization Transfer Ratio asymmetry percentage (MTR_asym_%), illustrates a significant and substantial CEST signal arising from the lactate·Eu-PCTA and lactate·Yb-PCTA ternary complexes, as illustrated in Figure 2D. This robust CEST effect observed with the ternary complexes sharply contrasts with the negligible CEST response observed with Eu-Yb-PCTA alone. As for Pr-PCTA, we hypothesize that the inability to detect a discernible CEST effect could stem from rapid or unusually slow proton exchange rates (*k*_*ex*_) with the bulk water. The *k*_*ex*_ for Yb- and Eu-PCTA, calculated using the Omega plot method at pH 6 and 7,^26^ were generally slower when compared to the values previously reported for DO3A-based SRs. Interestingly, both SRs exhibited a noteworthy minimum *k*_*ex*_ at pH 6, suggesting a similar acid-based proton catalytic effect (Table 1). The lower *k*_*ex*_ allows the application of relatively modest saturation powers (B_1_ < 12μT) to detect the CEST effect. This represents a clear advantage for *in vivo* translational studies where lower saturation powers are generally preferred.

To better understand the strength of the interaction, binding affinities (*K*_*A*_) were determined by acquiring CEST images of phantom tubes containing 20 mM SRs and titrations of lactate concentration between 0 and 600 mM at pH 6-7. The amplitude of the CEST effect was subsequently analyzed using a theoretical binding model outlined by Zhang *et al*.^8^. We assumed a single binding site for lactate in the model. The analysis enabled the determination of the binding affinity (*K*_*A*_) and the maximum CEST effect at saturation (Fig. 3A-B). Our findings revealed that all *K*_*A*_ values were ≤ 10^2^ M^-1^, underscoring a modest interaction between lactate and the SRs (Table 1). These results are particularly significant for translating SRs into non-invasive in vivo metabolic imaging applications in both preclinical and clinical settings. The determined *K*_*A*_ values indicate that lactate remains abundant in the bloodstream without being depleted by the SRs, confirming the feasibility of using these complexes for imaging purposes.

**Figure 3.**
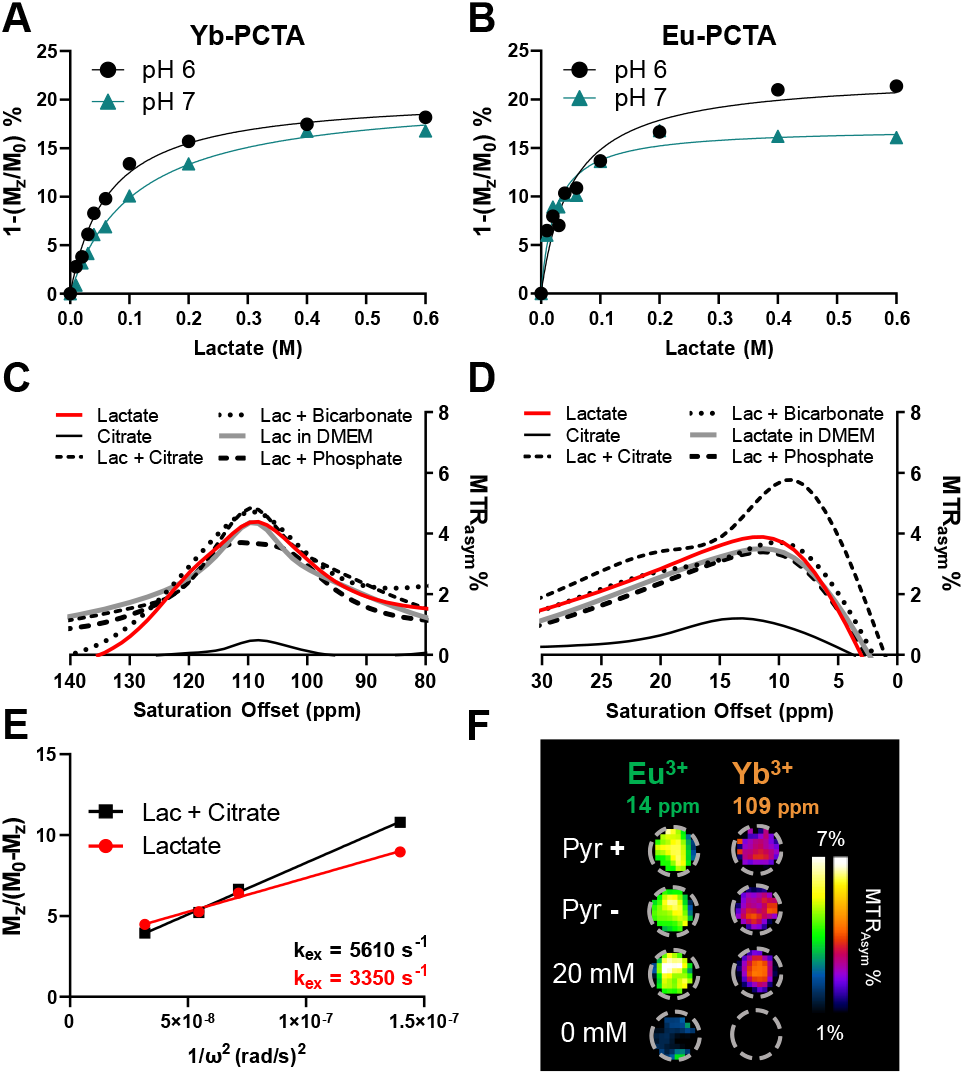
**A, B**. Effect of lactate titration (0-0.6 M) on the CEST effect produced by 20 mM Yb-PCTA (A) or Eu-PCTA (B) at pH 6 and 7. Lines = fitting line. CEST effect at 109 ppm (Yb-PCTA) and 14 ppm (Eu-PCTA) were obtained at 298 K after a pre-saturation of 21 µT, 5 s. **C, D**. MTR_asym_% plot of solutions containing 20 mM Yb-PCTA (C) or Eu-PCTA (D) and 40 mM lactate, citrate or different combinations of lactate and other metabolites and DMEM, pH 6, 298 K. **E**. Omega plot of Eu-PCTA with lactate or lactate and citrate (Lac + Citrate). **F**. CEST images of MC-38 colon adenocarcinoma cell growth media collected after 48h of culture with or without 1 mM pyruvate, mixed with 20 mM Yb-PCTA or Eu-PCTA.

The detection of lactate with the SRs also proved specific in the presence of potentially confounding -OH and -NH resonances present in cell growth media (DMEM). There were no notable differences in the CEST effect at 109 ppm when lactate-Yb-PCTA was combined with an equivalent amount of citrate, bicarbonate, or phosphate (Fig. 3C). In the case of Eu-PCTA, neither the addition of these metabolites nor the presence of DMEM significantly affected the CEST intensity at 10-14 ppm. The only exception was citrate, which notably generated a significantly higher CEST effect (Fig. 3D), but no CEST signal was observed for citrate-Eu-PCTA alone. The exchange rates for lactate-Eu-PCTA and citrate-lactate-Eu-PCTA were determined to be 3350 and 5610 s^-1^, respectively (pH 7, 21°C, Fig. 3E). This suggests that the difference in the CEST effect is not due to the citrate -OH CEST effect, but rather arises from a proton catalyzed buffering effect^27^. Recently, it has been reported that lanthanide complexes such as Eu-DO3A can also be used to detect inorganic phosphate.^28^ With the PCTA-based complexes, we did not observe any impact in the presence of 1 equivalent of phosphate. While it has been broadly demonstrated that phosphates interact with lanthanide-based contrast agents,^29^ inorganic phosphate levels vary from 1.1 to 1.2 mM in serum^30,31^ and may increase to 1.8-2.5 mM in the microenvironment of some tumors.^32,33^ Lactate, on the other hand, accumulates in the tumor microenvironment at usually higher concentrations (2-12.9 mM).^34,35^ Therefore, we do not foresee a significant impact of phosphate on detecting extracellular lactate in tumor tissue.

We also conducted lactate titrations in serum using both SRs. Results show a linear correlation between the CEST effect and lactate concentration (Fig. S2C), indicating that the SR offered a robust and specific tool for measuring lactate in biological solutions. According to the Human Metabolome Database (hmdb.ca), lactate is present in blood at concentrations 102-103 times higher than other metabolites such as pyruvate, malate, succinate, and citrate under physiological conditions. Therefore, we expect the impact of these molecules on the detection of extracellular lactate with CEST to be negligible. We hypothesize that the selective interaction between lactate and Ln-PCTA may be facilitated by its hydroxyl and carboxylate groups, which can effectively coordinate and stabilize the Ln-PCTA complexes.^36^ The geometry and electronic configuration of the metal ion within the complex, along with the spatial re-arrangement of the ligands, likely favor the selective binding of lactate over other monodentate anions (vide structural ^1^H NMR study in Fig. 2A and Fig. S4-7), as previously reported.^37,38^ Additionally, in pathological conditions such as cancer, the abundant presence of lactate likely enhances its preferential binding to Ln-PCTA.

To further test the potential of detecting extracellular lactate, we added 20 mM of the SRs to samples of MC-38 colon adenocarcinoma cell growth media collected after 48 hours of culture in a normoxic incubator. Furthermore, we cultured cells with or without pyruvate to test if the SRs could quantify small changes in lactate excretion. Pyruvate is the precursor of lactate in the glycolytic pathway and is usually provided in the growing medium of fast-growing cancer cells. We acquired CEST images of tubes containing the media, 20 mM lactate, or no lactate after titrating the pH to 6 in all samples (16 µT, 5-second saturation pulses as shown in Fig. 3F). Lactate concentration was calculated using calibration lines based on the CEST effect at 14 and 109 ppm.

To further explore the feasibility of *in vivo* CEST imaging of extracellular lactate with the SRs, we tested the cytotoxicity of the SR *in vitro* and compared it with that of a Gd-based contrast agent (Magnevist) and Eu-DO3A. No significant differences in cytotoxicity between these complexes were observed, even at the high concentrations required for CEST imaging (millimolar range) (Fig. S8).

Finally, we conducted an *in vivo* study using Yb-PCTA. We injected 0.2 mmol/Kg of a 1:1 mixture of lactate-Yb-PCTA intravenously into 3 healthy C57BL/6 mice. Then, we acquired CEST images at a fixed saturation offset (109 ppm) to dynamically monitor the bladder and muscle tissue over 60 minutes, as reported previously.^39,40^ We observed a CEST effect, represented as M_z_ magnetization difference compared to the baseline, in the bladder 20 minutes after injection, reaching a maximum at 50 minutes (Fig. 4A-C). This effect was not observed when we injected the same animals with Yb-PCTA dissolved in a 0.9% NaCl solution (Fig. 4B). The animal study demonstrated the safe *in vivo* detection of the ternary complex lactate-Yb-PCTA. The dynamics of the CEST effect in the bladder indicated that the SR was cleared quickly via renal elimination, with nearly complete clearance at 50 minutes post-injection (Fig. 4C). Based on the moderate affinities of lactate to Yb-PCTA, we hypothesize that the CEST signal detected in the bladder is in equilibrium between the initial lactate provided and the lactate secreted from the tissues. Nevertheless, we could not determine whether the lactate-Yb-PCTA ternary complex was excreted intact or if the complex was reconstituted in the urine—an acknowledged limitation.

**Figure 4.**
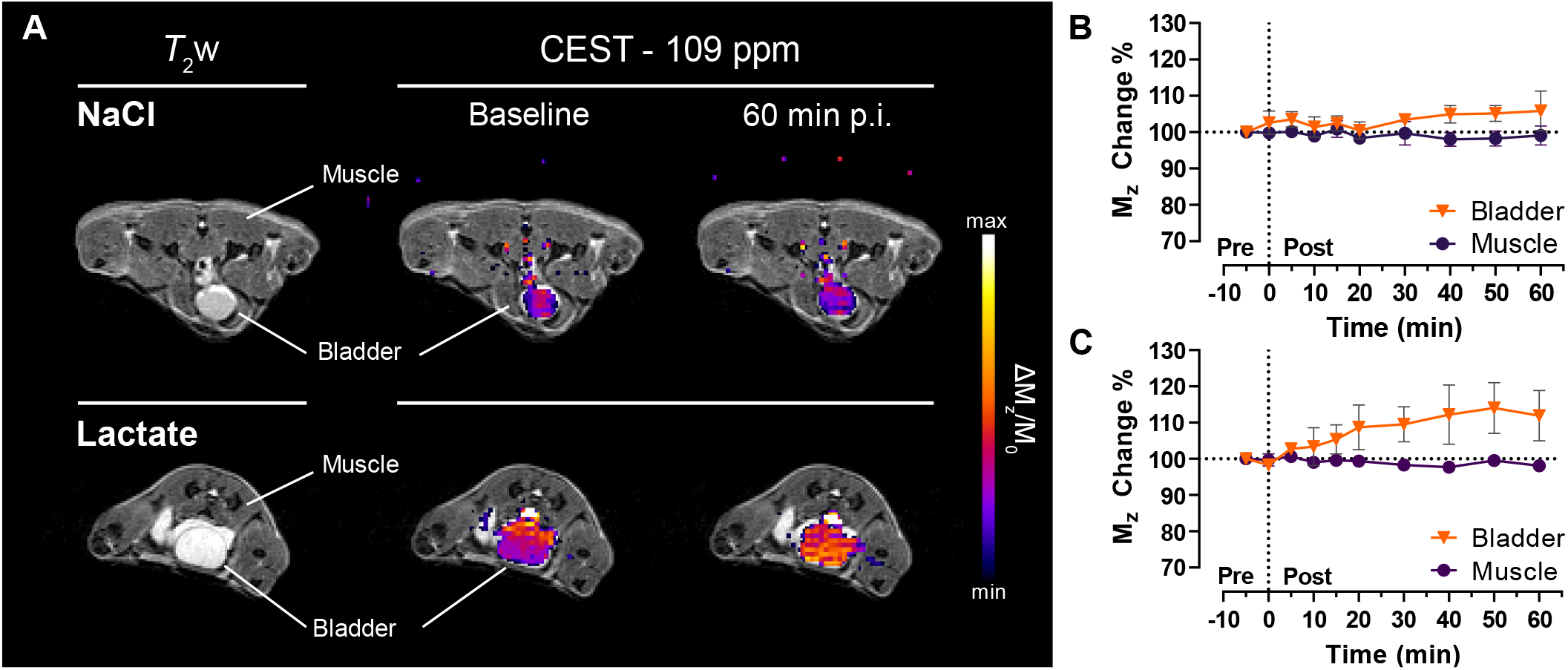
*In vivo* MR imaging of extracellular lactate with Yb-PCTA. **A**. Representative anatomical (*T*_2_-weighted) and extracellular lactate (CEST 109 ppm) images of healthy C57BL/6 mice before and 60 minutes after the intravenous injection of 0.2 mmol/Kg SR in NaCl solution or as a 1:1 mixture of lactate·Yb-PCTA. CEST images were acquired at a fixed offset (109 ppm) after 3 seconds of pre-saturation pulses of 14 µT. **B, C**. Quantification of the CEST effect in muscle tissue and bladder before (Pre) and after (Post) the injection of 0.2 mmol/Kg Yb-PCTA 1:1 lactate·Yb-PCTA (C) or in saline solution (B) (n = 3).

Our results demonstrate the feasibility of detecting lactate using CEST MRI with Ln-PCTA complexes; nevertheless, the *in vivo* validation is limited to a preclinical study with healthy mice. To evaluate its translational potential, further research is required to assess the efficacy of this approach in relevant disease models, such as tumor-bearing mouse models. Additionally, while the study demonstrates a robust CEST signal for lactate with Ln-PCTA complexes *in vitro*, the sensitivity of CEST MRI for detecting extracellular lactate *in vivo* may be influenced by factors such as tissue heterogeneity and delivery of the SR to the target tissues. Further ongoing work to improve the CEST signal-to-noise ratio will help achieve quantitative metabolic imaging.

## Conclusions

Our study showed that Ln-PCTA complexes can form ternary complexes with lactate. High-resolution NMR analysis successfully demonstrated the interaction between these complexes and lactate. Remarkably, Yb-PCTA and Eu-PCTA showed only a TSD isomer structure in solution at several temperatures when coordinated with lactate.^22,23^

We assessed the CEST effect and its specificity for lactate in various conditions. Yb-PCTA and Eu-PCTA exhibited strong and unique CEST signals in the presence of lactate, supporting their potential as effective CEST shift reagents. The *K*_A_ values indicated a moderate binding between lactate and the shift reagents, suggesting that physiological lactate levels remain stable in the bloodstream during the use of these SRs for *in vivo* molecular imaging. Our experiments in cell culture media and *in vitro* cytotoxicity assessments demonstrated that these SRs are safe and optimal for non-invasive *in vivo* imaging. Furthermore, the *in vivo* detection of the lactate-Yb-PCTA ternary complex validated the safety and efficacy of this approach. Our experiment demonstrated that a 0.2 mmol/Kg dose of Yb-PCTA—similar to the clinical dose of Gd-based contrast agents and much lower than typical iodine CT agents—could be detected intact in the mouse bladder, where most of the injected dose is excreted. While detecting extracellular lactate in tumors or organs outside the main excretion routes may require higher SR doses, ongoing optimization of the CEST acquisitions, the *k*_*ex*_ and *K*_*A*_ of lactate-Ln-PCTA complexes may compensate for the moderate CEST SNR generated. This optimization could potentially enhance the CEST effect, allowing for lower injection doses and weaker pre-saturation pulses for CEST detection. The findings of this study provide valuable insights for developing non-invasive metabolic imaging probes in preclinical and clinical settings. This study demonstrates the potential of combining CEST MRI with safe inorganic lanthanide-based PCTA complexes to profile various cancer cell lines and the tumor microenvironment effectively.

## Supporting information

ESI file

## ABBREVIATIONS

CEST: chemical exchange saturation transfer
DMEM: Dul-becco’s Modified Eagle Medium
DOTA: 1,4,7,10-tetraazacy-clododecane-1,4,7,10-tetraacetic acid
DO3A: 1,4,7,10-tetraazacyclododecane-1,4,7-triacetic acid
MRI: magnetic res-onance imaging
MTR_asym_%: Matgnetization Transfer Ratio asymmetry percentage
NMR: Nuclear Magnetic Resonance
SAP: square antiprism
SR: shift reagent
PCTA: 3,6,9,15-tetraazabicyclo[9.3.1]pentadeca-1(15),11,13-triene-3,6,9-tri-acetic acid
TSAP: twisted square antiprism
TSD: twisted snub disphenoid.

## ASSOCIATED CONTENT

Detailed descriptions of all the experimental procedures, complex synthesis, cytotoxicity results, and additional relevant ^1^H NMR and CEST spectra are listed in Supplementary Information. This material is available free of charge via the Internet at http://pubs.acs.org.

## AUTHOR INFORMATION

## Author Contributions

The manuscript was written with contributions from all authors. All authors have approved the final version of the manuscript.

## ACKNOWLEDGMENT

A.F.M acknowledges the support from the the Deutsche Forschungsgemeinschaft (DFG, German Research Foundation) – 516238665 and 527345502. This work was supported by the Cluster of Excellence iFIT (EXC 2180) “Image-Guided and Functionally Instructed Tumor Therapies,” University of Tuebingen, Germany, the Werner Siemens Foundation, the Alexander von Humboldt Foundation within the framework of the Sofja Kovalevskaja Award (to AFM), and the DKTK German Cancer Consortium Innovation program “HYPERBOLIC. C.F.G.C.G and P.F.C. acknowledge the Foundation for Science and Technology (FCT), Portugal, for funding the CQC-IMS (UID/QUI/00313/2020, UIDP/00313/2020 and POCI-01-0145-FEDER-027996) of the University of Coimbra.

