## Supplementary material for "Metabolic PCTA-Based Shift Reagents for the Selective Detection of Extracellular Lactate Using CEST MRI": ESI file

#### Supplementary information

### Chemicals and synthesis

Pyclen was dissolved in acetonitrile. Potassium carbonate was added as a base. The solution was cooled in an ice bath before adding t-butylbromoacetate. The solution was allowed to warm to room temperature and then stirred overnight. The solution was filtered to remove the salts and then split into two separate vessels. An equivolume of water was added to each. 1N HCl was added to lower the pH to 3-4. An equivalent of 0.5M Europium chloride, Ytterbium chloride, or Praseodymium chloride was added. The Eu & Yb solutions were heated to 45-55°C for complexation. The Pr solutions were heated to 75°C for complexation. Throughout the day to form the complex, 1N NaOH was used to maintain a pH = 4-6 ($Eu^{3+}$), pH = 5.3-5.6 ($Yb^{3+}$), and pH = 5.8-6.5 (Pr). Acetonitrile was replenished as needed to maintain a consistent volume. The complexation was monitored by HPLC. After complexation, the metal acts as a catalyst to remove the t-butyl esters. The reactions were stopped after > 80% completion. The solutions were used as a stock solution for preparative HPLC purification. A Phenomenex Luna C18(2) column connected to a Waters DeltaPrep system was used. A simple gradient using 0.025% TFA in $CH_3CN/H_2O$ modifiers was used. Collected fractions were freeze-dried to obtain the purified complexes. Final purities were $\geq$ 98% by HPLC. Identities were verified by mass spectrometry. The solids were analysed by ICP-MS to quantify the metal content: 21.8% Eu in Eu-PCTA,

24.9% Yb in Yb-PCTA, and 17.4% Pr in Pr-PCTA. Based on the percent metal values, it is likely the compounds were isolated as a 1TFA salt with a small percentage of residual water. Isolation of a 1TFA salt has been observed in our lab with similar complexes. Percent yields are based on the formula weight of the 1TFA salt. Chemical structures and IUPAC names were obtained using Chemaxon MarvinSketch 24.1.2.[1]

Table S1.

| Complex | Metal Content | Neutral Molecule Fw: % Metal | TFA salt Fw: % Metal | Yield |
|---|---|---|---|---|
| Eu-PCTA | 21.8% | 529.3 g/mol: 28.7% | 643.4 g/mol: 23.6% | 56% |
| Yb-PCTA | 24.9% | 550.4 g/mol: 31.4% | 664.4 g/mol: 26.0% | 48% |
| Pr-PCTA | 17.4% | 518.3 g/mol: 27.2% | 632.4 g/mol: 22.2% | 22% |

#### CEST MRI Experiments

Phantoms were acquired using a 7 T (300 MHz) preclinical MRI scanner (Bruker BioSpec 70/30, Bruker BioSpin, Ettlingen, Germany) using an 86-mm diameter $^1$H transceiver volume coil (Bruker). 2D CEST coronal images were acquired using a previously reported FISP sequence with the following parameters:[2] echo time (TE) 1.80, repetition time (TR) 3.60, flip angle 30°, field of view (FOV) 100x80 mm, slice thickness 1 mm, matrix size 128x80, resolution 0.78x0.75x1 mm. CEST pre-saturation consisted of 5 seconds, continuous pulses with a $B_1$ ranging between 2 and 21 µT depending on the experiment.

#### Z-spectra

Z-spectra were acquired in phantom tubes containing 50 mM of Yb- and Eu-PCTA (Pr-PCTA: 40 mM) mixed with 50 mM lactate (for Pr-PCTA: 40 mM) in 50 mM HEPES buffer at pH 6 and 7, 298 K. 5 seconds pre-saturation pulses were used with different $B_1$ values depending on the experiment ($B_1$: 8, 12, 20 µT, Figure S1; $B_1$: 16 µT, Figure 2-3). Raw spectra were fitted to Lorentzian line shapes based on a two-pool model (water, lactate·Ln-PCTA) using an in-house written MATLAB script. CEST effect with respect to saturation offset (Δω) was quantified as Magnetization Transfer Ratio asymmetry percentage ($MTR_{asym}$%), as per $MTR_{asym}\% = [(M_z^{-\Delta\omega} - M_z^{+\Delta\omega})/M_0] \times 100$, unless differently specified.

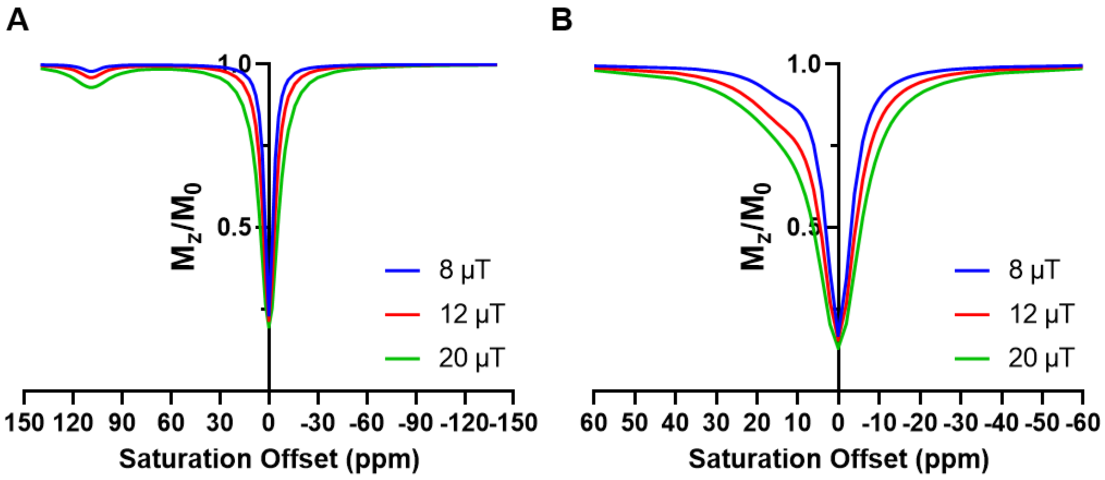

Figure S1. Z-spectra of 50 mM Yb-PCTA (A) or Eu-PCTA (B) mixed with 50 mM lactate acquired at 298 K, pH 6.9, with different $B_1$ values.

#### Calibration curves for lactate determination with CEST MRI

Phantoms containing 20 mM SRs mixed with 0-40 mM lactate in 50 mM HEPES buffer or human serum (Sigma-Aldrich) at pH 6 and 7, 298 K, were used to generate calibration curves by obtaining linear regression lines between the amplitude of CEST effect at 14 ppm (Eu-PCTA) or 109 ppm (Yb-PCTA) and lactate concentration (Figure S2). CEST pre-saturation consisted of 5 seconds, 16 µT continuous pulses. Z-spectra were fitted to Lorentzian line shapes (two pools, water, and lactate·Ln-PCTA), and then the amplitude of the CEST effect (CEST%) was calculated with an in-house written MATLAB script. CEST images were acquired in triplicate.

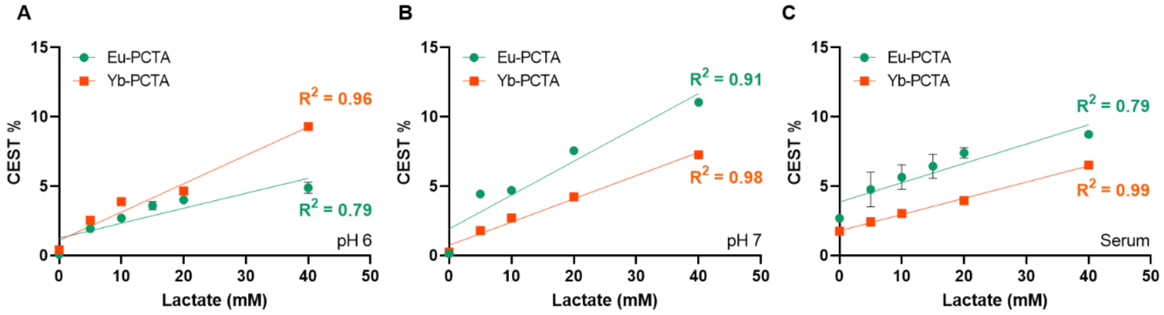

Figure S2. Plot of CEST effect at 14 ppm (Eu-PCTA) or 109 ppm (Yb-PCTA) vs lactate concentration, at pH 6 (A), pH 7 (B), or human serum at pH 7 (C). Goodness of the fit ($R^2$) of the linear regression analysis is reported for each set of lactate titration. Data expressed as mean CEST % ± SD.

#### Exchange rates determination

Exchange rates were determined using the Omega plot method.[3] 20 mM, 1:1 solution of lactate and SRs at pH 6 or 7 were scanned as described before using CEST pre-saturation pulses with $B_1$ ranging between 2 and 20 µT.

#### Binding affinity determination

Binding affinity for the SRs-Lactate complexes was determined using the $CEST_{\%}$ effect at 14 ppm (Eu-PCTA) or 109 ppm (Yb-PCTA) at pH 6 and 7, 298 K. $CEST_{\%}$ was determined as per CEST% = $[1-(M_z^{\Delta\omega}/M_0)] \times 100$. CEST images were acquired after a pre-saturation pulse of 5 seconds, 21 µT $B_1$, using 20 mM SRs and lactate concentrations ranging between 0 and 600 mM. After background correction, $CEST_{\%}$ for each lactate concentration was fitted according to the following equation to determine $K_a$ and $CEST_{\%}$ at saturation ($CEST_{bound}$):[3]

$$CEST_{\%} = \frac{CEST_{bound} \cdot \left(NC_{EuDO3A} \cdot C_{EuDO3A} + C_{Lactate} + K_a^{-1} - \sqrt[2]{(NC_{EuDO3A} \cdot C_{EuDO3A} + C_{Lactate} + K_a^{-1})^2 - 4 \cdot NC_{EuDO3A} \cdot C_{EuDO3A} + C_{Lactate}}\right)}{2 \cdot C_{EuDO3A}}$$

#### Competition experiment

Selectivity of the CEST effect for lactate was tested in phantoms containing 20 mM of SRs and different combinations of 40 mM lactate, citrate, $NaHCO_3$, $NaH_2PO_4$, Dulbecco's Modified Eagle's medium (DMEM) at pH 7, 298 K. CEST images were acquired as described above with 5 seconds pre-saturation pulses with a $B_1$ of 16 µT. Exchange rates were determined with the Omega plot as described in "Exchange rates determination."

#### *In vivo* ShiftCEST

Animal experiments with healthy C57Bl/6 mice were conducted following German federal regulations on the use and care of experimental animals and approved by the local authorities

(Regierungspräsidium Tübingen). 3 healthy female mice were anesthetized using 1.5% isoflurane in pure oxygen, and a catheter was placed in a tail vein. Axial 2D CEST images were acquired with the same FISP sequence used for phantoms, with the following parameters: TE 1.76 ms, TR 3.52, flip angle 30°, FOV 38.4x38.4 mm, slice thickness, matrix 64x64, spatial resolution 0.6x0.6x1 mm. 20 pre-injection CEST images were acquired at 109 ppm after a pre-saturation of 3 seconds, 14 µT continuous pulses to determine the baseline, then further CEST images were acquired every 5 minutes for the first 30 minutes and at 40, 50 and 60 minutes after i.v. injection of 0.2 mmol/Kg of Yb-PCTA dissolved in 0.9% saline solution. After 48 h, the experiment was repeated by injecting a 1:1 mixture of 0.2 mmol/Kg lactate*Yb-PCTA. 20 CEST images were acquired at each time point and averaged. The dynamics of the CEST effect were determined for bladder and muscle tissue as $[1-(M_z{}^{\Delta\omega}{}_{(t)}/M_z{}^{\Delta\omega}{}_{(\text{pre-injection})})] \times 100$.

Anatomical $T_1$ weighted- (FLASH, TE 2.67 ms, TR 8.85 ms, flip angle 10°, matrix 272x136x96, FOV 76.8x34.8x22.8 mm, resolution 0.3x0.3x0.3 mm) and $T_2$ weighted- (RARE, TE 90.51 ms, TR 1800 ms, flip angle 90°, matrix 256x116x76, FOV 75x35x21.12 mm, resolution 0.28x0.26x0.22 mm) images were also acquired before and 60 minutes after injection.

#### Relaxometry experiments

To determine $r_1$ and $r_2$ relaxivity, 0.1-1 mM Eu-PCTA phantoms and 0-10 mM Yb-PCTA, dissolved in 50 mM HEPES buffer or human serum, were prepared in 0.3 mL tubes. $T_1$-weighted MR images of phantoms were acquired using a 2D T1-FLASH sequence with the following parameters: TE 4 ms, TR 100 ms, flip angle 50°, field of view 50x50 mm, 10 slices, slice thickness 1 mm, matrix size 192x192, resolution 0.260x0.260x1 mm. $T_1$ maps were acquired using a standard 2D RARE VTR sequence with 15 TRs: 50, 100, 200, 300, 400, 500, 600, 700, 800, 900, 1000, 1500, 2500, and 3000 ms. Other parameters: TE 8 ms, field of view 50x50 mm, 2 slices, slice thickness 1 mm, matrix size 192x192, resolution 0.260x0.260x1 $mm^3$. $T_2$-weighted MR images were acquired using a 2D T2-TurboRARE sequence with the following parameters: TE 28 ms, TR 1800 ms, flip angle 90°, field of view 50x50 mm, 20 slices, slice thickness 1 mm, matrix size 256x192, resolution 0.195x0.260x1 mm. $T_2$ maps were acquired using a standard 2D MSME sequence with 60 TEs ranging from 8 to 480 ms. Other parameters: TR 3000 ms, field of view 50x50 mm, 5 slices, slice thickness 1 mm, matrix size 192x192, resolution 0.260x0.260x1 mm. Experiments were performed at 298 K in Tris buffer and 298 and 310 K in human serum. $T_1$ and $T_2$ maps were generated using the MRI Analysis Calculator for ImageJ.

#### NMR Experiments

Nuclear magnetic resonance (NMR) solutions were prepared using deuterated solvent $D_2O$. The $^1H$ NMR spectra were acquired on a Bruker Avance III 400 spectrometer (Bruker, Massachusetts, USA) operating at a frequency of 400.13 MHz ($^1H$), at various temperatures, using a 5-mm z-gradient inverse probe. For 1D data acquisition, the *zgpr* pulse sequence was implemented with 128 k complex points, a spectral width of 200000 Hz, a recycle delay of 0.05 s, and a 90° pulse width of 13.17 µs and 1024 transients (Figures S4-S6). 2D COSY and EXSY spectra were acquired on Bruker Avance NEO 600 spectrometer, using BBFO 5 mm iProbe (Figure S7). Subsequently, the resulting data was processed and analyzed using Topspin v4.0 (Bruker) and MestReNova 9.1 (Mestrelab).

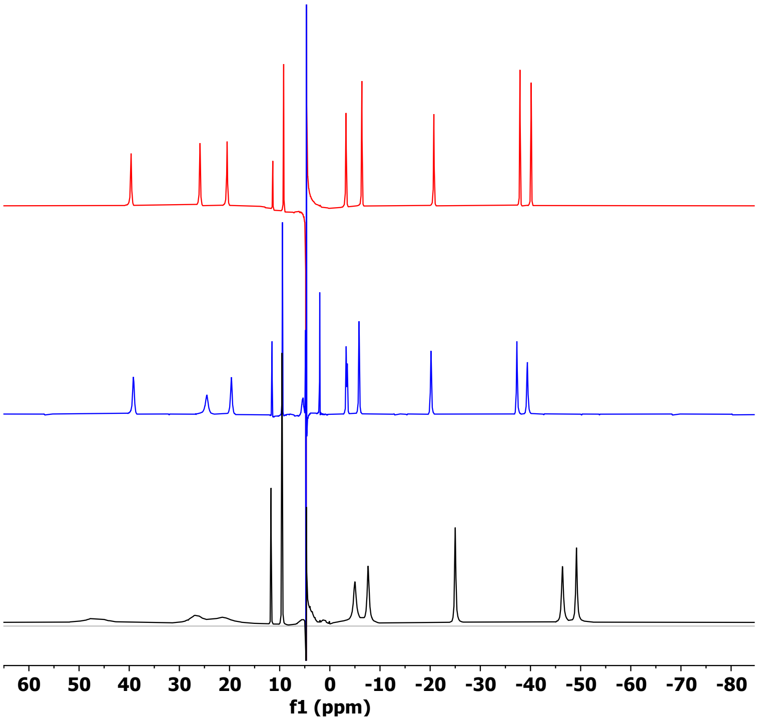

Figure S3. $^1$H NMR spectra of Yb-PCTA at 298 K (black), 2:1 Lactate: Yb-PCTA mixture at 363 K (blue) and Yb-PCTA at 363 K (red)

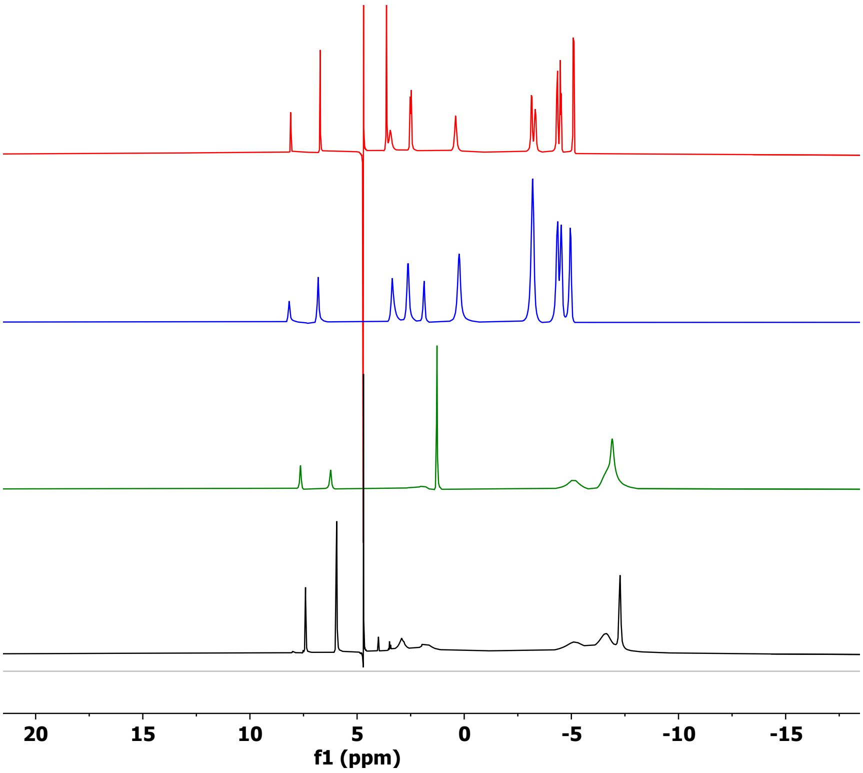

Figure S5. $^1$H NMR spectra of Eu-PCTA at 298 K (black), 2:1 Lactate: Eu-PCTA mixture at 298 K (green), 2:1 Lactate: Eu-PCTA mixture at 363 K (blue) and Eu-PCTA at 363 K (red).

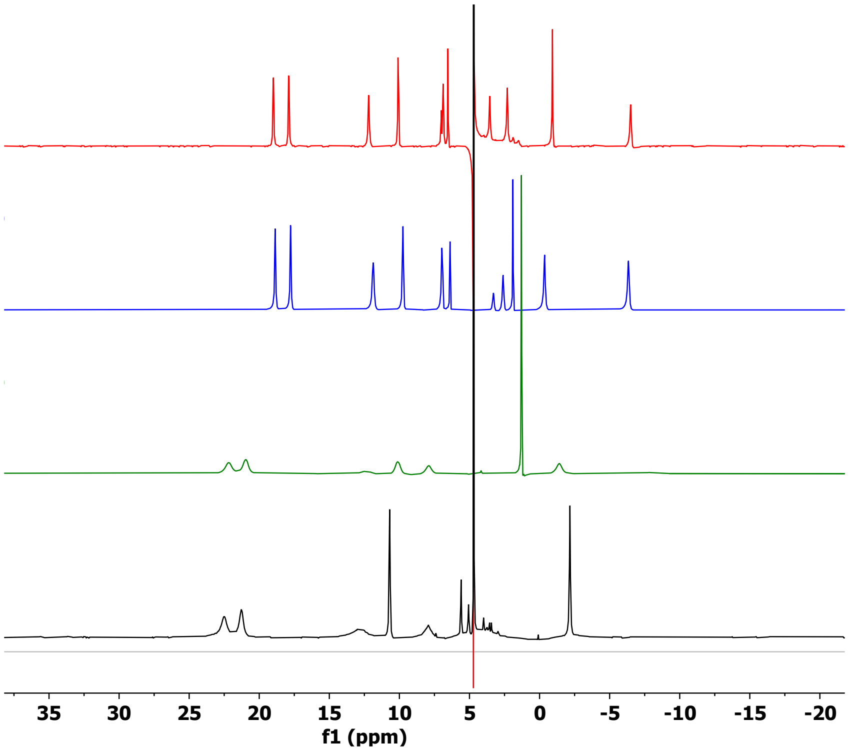

Figure S6. $^{1}H$ NMR spectra of Pr-PCTA at 298 K (black), 2:1 Lactate: Pr-PCTA mixture at 298 K (green), 2:1 Lactate: Pr-PCTA mixture at 363 K (blue) and Pr-PCTA at 363 K (red).

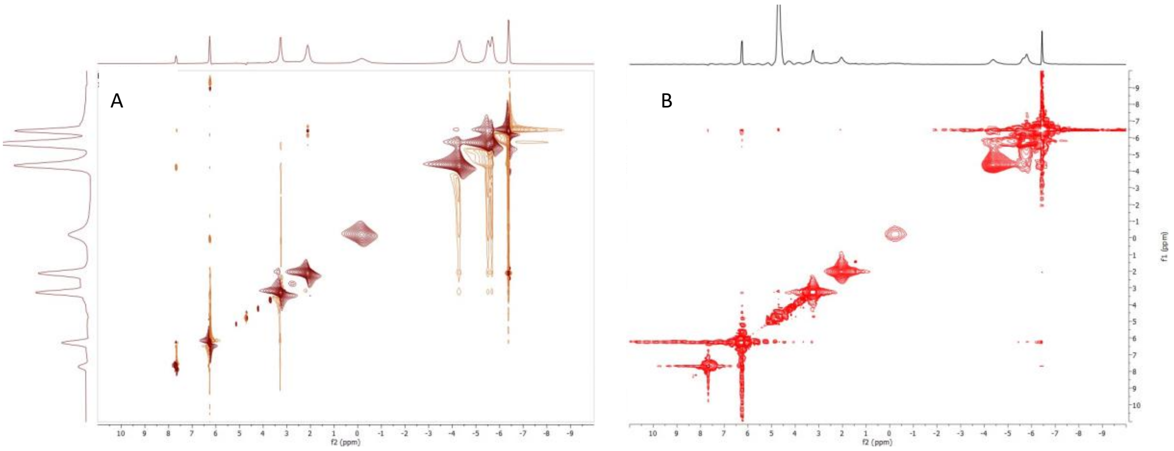

Figure S7. A. 2D-COSY spectrum of an aqueous solution of Eu-PCTA at 298 K (pH 7.1), B. 2D-EXSY spectrum of an aqueous solution of Eu-PCTA at 298 K (pH 7.1; mixing time, 10 ms).

#### Cytotoxicity

Cytotoxicity of Yb-, Eu-, and Pr-PCTA complexes was evaluated using an MTS assay (Promega, Madison, WI, USA) and compared to the previously reported shift reagent (Eu-DO3A). MC-38 cells (Kerafast) were cultured in T175 flasks with Dulbecco's Modified Eagle's Medium (DMEM) supplemented with 10% FCS, 1% penicillin/streptomycin, L-glutamine, sodium pyruvate, and 15 mM HEPES in a humidified incubator (5% $CO_2$, 37°C). For experiments, cells were trypsinized, counted with Trypan Blue, and seeded at a density of 2000 cells/well in 96-well plates. After 24 h, cells were incubated for 2 h with 5 mM Yb-, Eu-, Pr-PCTA, 5 mM Eu-DO3A or 10 µM Magnevist (Bayern) dissolved in phenol red-free DMEM, then the MTS reagent mix was added. After further incubation at 37°C for 2 or 4 h, absorbance at 490 nm was measured. Cytotoxicity was determined as viability versus control by comparing the absorbance of treated cells to control cells, expressed as %. Cytotoxicity of Ln-PCTA complexes was pooled and compared with that of Eu-DO3A and Magnevist (Figure S8).

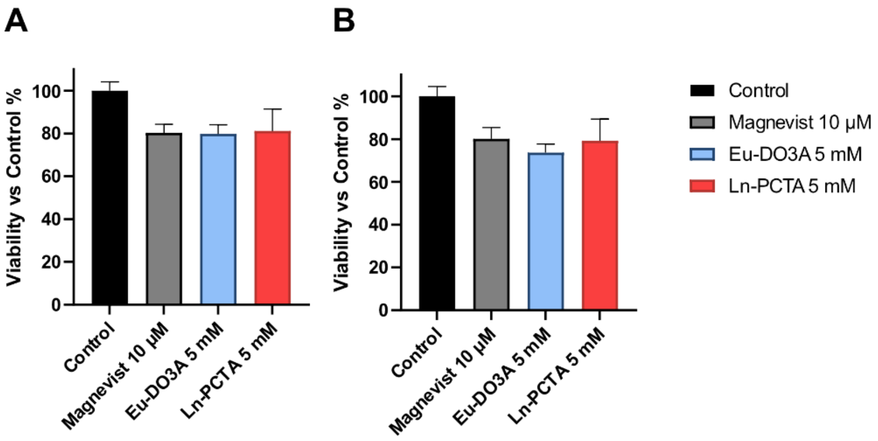

Figure S8. Cytotoxicity of Ln-PCTA complexes compared to Magnevist and Eu-DO3A. Cytotoxicity was determined with an MTS assay and calculated as percentage of viability compared to control cells (non-treated cells) after 2h (**A**) or 4h (**B**) of incubation. Data expressed as mean ± SEM.

#### Cell culture experiment to detect lactate excreted by cancer cells

MC-38 cells were cultured in T175 flasks with DMEM supplemented with 10% FCS, 1% penicillin/streptomycin, L-glutamine, sodium pyruvate, and 15 mM HEPES in a humidified incubator (5% $CO_2$, 37°C). For experiments, cells were trypsinized, counted with Trypan Blue, and seeded at a density of $5\times10^5$/well in 6 well plates in a medium supplemented with or without 1 mM sodium pyruvate. After 48h, the growing medium was collected, centrifuged for 30 minutes at 4°C in 10 kDa filter tubes, and stored at -80°C until further use, while cells were harvested and counted with Tripan blue. For CEST measurements of lactate excreted by the cancer cells, the filtered growing media were transferred to 0.3 mL tubes, Yb- or Eu-PCTA was added to a final concentration of 20 mM, and pH was adjusted to 7.0. CEST images were acquired as described before, using pre-saturation pulses of 5 seconds with a $B_1$ of 16 µT. Lactate-CEST ($1-M_z/M_0$ %) calibration lines were used to quantify lactate concentration. An enzymatic lactate dehydrogenase (LDH) kit (Lactate Assay Kit II, Sigma-Aldrich) was used to cross-validate lactate concentration. Lactate concentrations determined via the enzymatic kit and the CEST images are reported in Table S2.

Table S2. Lactate concentrations were determined with the LDH kit and CEST MRI with Yb- and Eu-PCTA in samples of MC-38 cells culturing medium, expressed in mM. Cells were cultured for 48h in the absence (Pyr -) or presence 1 mM pyruvate (Pyr +). The CEST effect at 14 ppm (Eu-PCTA CEST) or 109 ppm (Yb-PCTA CEST) was used to calculate lactate concentration thanks to a lactate-CEST calibration line.

| Lactate (mM) | Pyr + | Pyr - |
|---|---|---|
| LDH Kit | 24.6 | 22.8 |
| Yb-PCTA CEST | 25.1 | 22.2 |
| Eu-PCTA CEST | 24.8 | 19.9 |
